# Transcriptomic Dissection of the AMF-*Solanum melongena* Interaction: Unveiling Molecular Secrets

**DOI:** 10.1101/2024.02.17.580814

**Authors:** Subhesh Saurabh Jha, L.S. Songachan

## Abstract

Fungus-based biofertilizers, in particular, have been shown to improve plant growth and health by providing essential nutrients and improving soil health. Arbuscular mycorrhizal fungi (AMF) form mutualistic symbiotic relationships with plants, including those in the Solanaceae family. They colonize the root systems of plants and aid in enhancing nutrient absorption efficiency, particularly in phosphorus-deficient soils. The present study was carried out to study the interaction between *Solanum melongena* L. with *Glomus macrocarpum* and *Funneliformis mosseae* (formely known as *Glomus mosseae*) at transcript level. In this study a total of 365 transcripts were upregulated (>1.5x) folds in *S. melongena* in response to both the fungi, while 44 transcripts were upregulated only in response to *G. macrocarpum* and 28 transcripts were upregulated only in response to *F. mosseae*. Similarly, 49 transcripts were downregulated less than −1.5 folds in response to both the fungi while 78 were downregulated only in response to the *G. macrocarpum* and 36 were downregulated only in response to *F. mosseae*.

KEGG pathway analysis of *S. melongena* treated with *G. macrocarpum* revealed carbon metabolism, cofactor biosynthesis and Endocytosis as the dominant metabolic pathway, while analysis of the *F. mosseae* treatment revealed Glycerophospholipid and Endocytosis metabolism as dominant metabolic pathways.

## Introduction

Plants play a significant role in our ecosystem and human survival, and agriculture is the cornerstone of global food security. Investigations into the environmentally-friendly and sustainable methods are needed to boost crop yields in order to meet the growing global demand for food.

In the recent years, studies on the role of microorganisms in plant growth and sustainable agriculture has increased. Fungus-based biofertilizers, in particular, have been shown to improve plant growth and health by providing essential nutrients and improving soil health. Particularly, the advantageous effects of arbuscular mycorrhizal fungi (AMF) on a variety of crops, such as cereals, legumes, and vegetables, have been extensively studied (Chen et al., 2018; Begum et al., 2019; Khan et al., 2022; Sui et al., 2022).

Most AMF species are members of the phylum Mucoromycota’s sub-phylum Glomeromycotina. (Spatafora et al., 2016). This sub-phylum contains 25 species and four orders of AMF, namely Glomerales, Archaeosporales, Paraglomerales, and Diversisporales (Redecker et al., 2013). They use lipids and products from plant photosynthetic processes to complete their life cycle since they are obligatory biotrophs (Bago et al., 2000, Jiang et al., 2017).

*Glomus macrocarpum* and *Funneliformis mosseae* are both species of arbuscular mycorrhizal fungi (AMF) that form mutualistic symbiotic relationships with plants, including those in the Solanaceae family. They colonize the root systems of plants and aid in enhancing nutrient absorption efficiency, particularly in phosphorus-deficient soils (Aziz et al., 2011). These fungi develop structures in the root cells of plants called arbuscules, which expand the root system’s surface area and enable a direct interchange of nutrients between the fungus and the plant. This causes the plant to absorb more nutrients, which improves plant development and productivity.

The vegetable crop brinjal (*Solanum melongena* L.), which is widely grown in tropical and subtropical regions, may be a potential candidate for the use of fungus-based biofertilizers. The impact of these biofertilizers on the proteins in brinjal is not well understood, though. More effective and environmentally friendly agricultural practices might come from a better understanding of the molecular mechanisms by which fungus-based biofertilizers interact with and alter the physiology of brinjal.

Systems biology is a cross-disciplinary area of research that brings together numerous fields, including biology, mathematics, computer science, and engineering, to comprehend complex biological systems. In order to develop a thorough understanding of biological processes, the system biology entails the integration of high-throughput experimental data with computer models.

Systems biology approaches have fundamentally changed how we understand the way plants grow and develop as well as the way they react to environmental stress. Plant systems biology is the study of the structure, function, and interactions of plant molecules and networks using high-throughput technologies. These technologies include genomics, transcriptomics, proteomics, and metabolomics. The system biology has become a potent tool for comprehending intricate biological processes in plants, and it is anticipated to be crucial in the advancement of sustainable agriculture

The purpose of the research is to employ the system biology to understand the mechanism via which the fungal biofertilizers affect the transcriptome of brinjal. The investigation would clarify the molecular mechanisms and pathways underpinning the effects of fungus-based biofertilizers on brinjal.

## Materials and Methods

### Study site and Sampling

The study was conducted at botanical garden, department of botany, Banaras Hindu University, Varanasi, India located at 25° 16’ 4.3608’’ N and 82° 59’ 25.7784’’ E with an altitude of 81 metres above the sea level.

### Inoculation of *Solanum melongena* L. plants

For the purpose, pot culture experiment was set up. Control and both the treatments were set up in triplicates. Pure culture of Funneliformis mosseae and Glomus macrocarpum was obtained by monoculturing both these isolates of AMF. For biofertilizer experiment brinjal seeds disinfected with 1% sodium hypochloride was placed on sterilized sand-soil substrate(1:1v/v) in 200 ml disposable plastic containers. The germination set up were kept in B.O.D incubator at 25[under white fluorescent tubes (Photoperiod 12 hr) and watered whenever required to keep the soil mixture moist. After one month, germinated seedlings of S.melongena were transferred in 8 x 7.6 inch clay pots containing sterilized sand-soil substrate(1:1 v/v) with 10g each of both the inoculants. Uninoculated(control) plants were also maintained. AMF colonisation was ascertained by advanced microscopy.

Transcriptomics studies was done for the inoculations.

## Results

The paired-end libraries of the transcripts of *Solanum melongena* L. plant treated without or with arbuscular mycorrhizal fungi, *Glomus macrocarpum* and *Funneliformis mosseae* were sequenced on the Illumina HiSeq platform.

**Figure 1.**
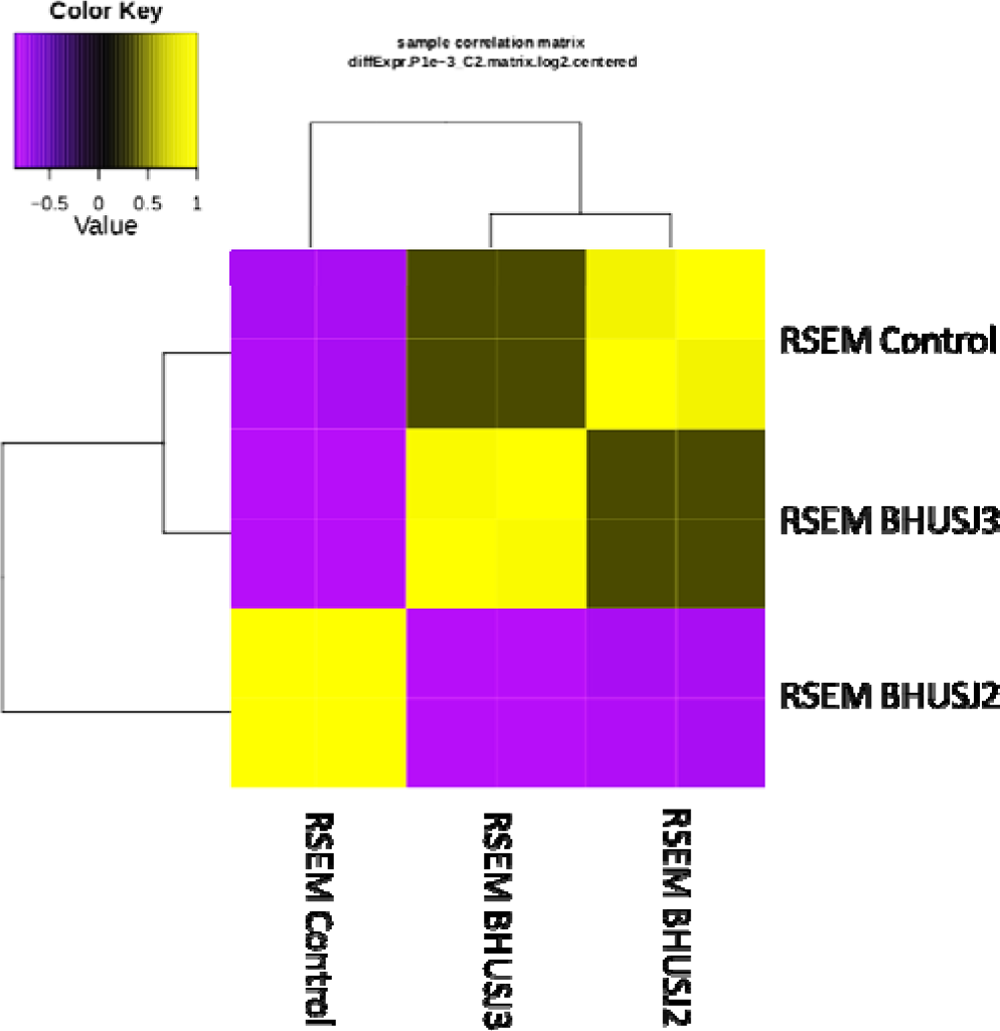
Sample Correlation Matrix of the transcriptome libraries

The paired-end libraries of the transcripts of *Solanum melongena* L. plant treated without or with arbuscular mycorrhizal fungi, *G. macrocarpum* and *F. mosseae* were sequenced on the Illumina HiSeq platform. The control samples i.e., *S. melongena* L. plant not treated with the AMF had mean number of 21722156.5 reads (78.81%) with 109050 fragments per kilobase per million mapped reads (FPKM) value greater than or equal to 1.0. Similarly, BHUSJ2 sample i.e., *S. melongena* L. plant treated with *Glomus macrocarpum* had mean number of 2150827 reads (80.65%) with 57733 FPKM value ≥ 1.0 and BHUSJ3 sample i.e., *S. melongena* plant treated with *Funneliformis mossae* had mean number of 25187514 reads (81.52%) with 95655 FPKM value ≥ 1.0.

The sample correlation matrix of the transcriptome libraries revealed that BHUSJ3 sample was closer to the control samples as compared to the BHUSJ2 samples (Figure 2). The observation implies that the *Glomus macrocarpum* had more pronounced effect on the transcriptomics of the *S. melongena* than that of the *Funneliformis mossae*.

**Figure 2.**
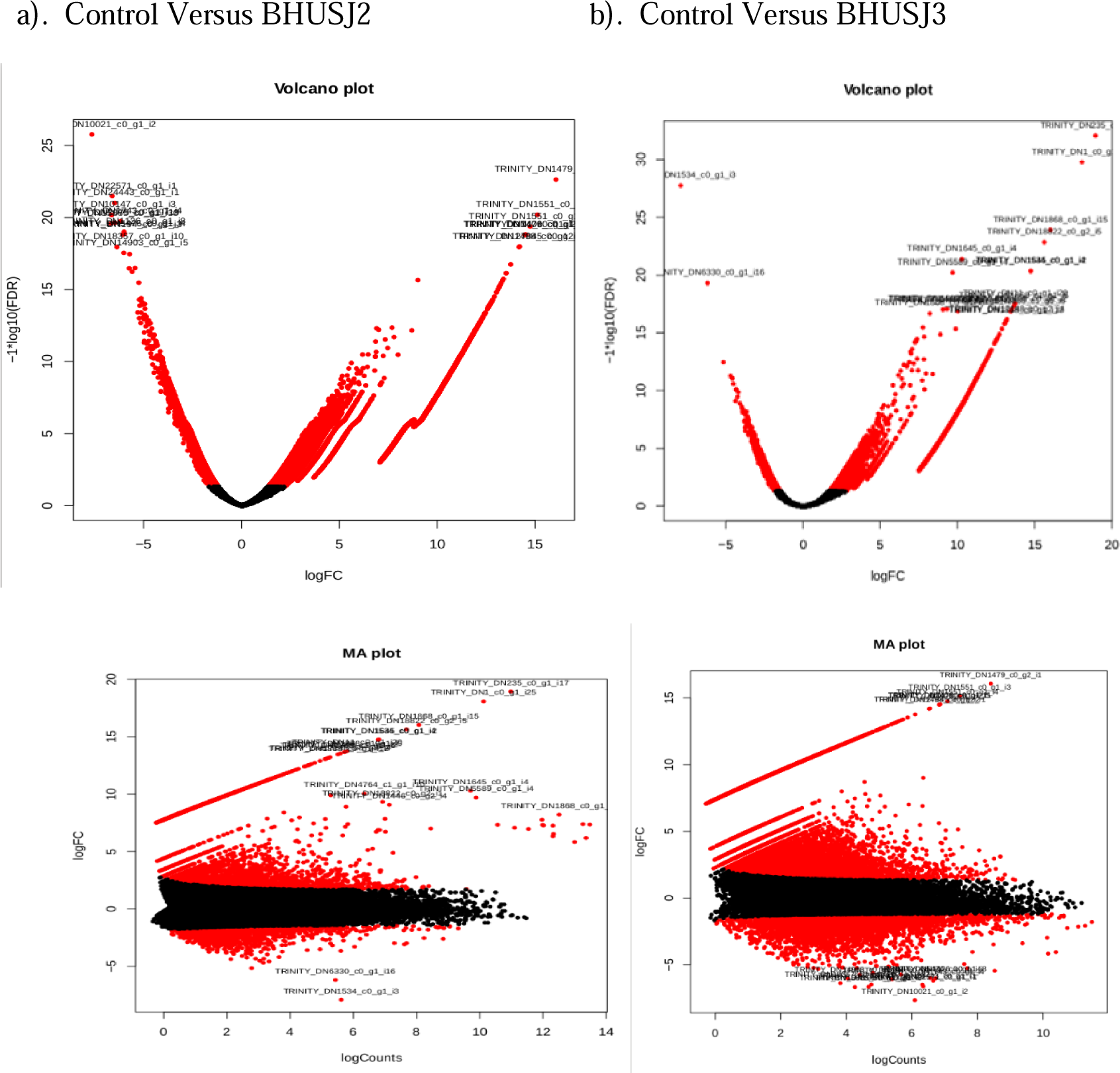
MA and Volcano plots of the differentially expressing libraries

The volcano plots and MA plots of the differentially expressed genes (DEGs) of the libraries are shown in the figure 3. Comparisons of the control library with BHUSJ2 i.e., genes of *S. melongena* when treated with the *Glomus macrocarpum*, revealed nearly 56897 DEGs, of which 33136 were statistically significant (Figure 3a). Out of the differentially expressed genes included 7608 upregulated transcripts and 25528 downregulated transcripts. of the upregulated transcripts 2685 genes were upregulated more than 1.5 folds while among the downregulated transcripts, 1285 genes were downregulated below −1.5 folds. Similarly, the comparisons of the control library with BHUSJ3 library i.e., genes of *S. melongena* when treated with the *Funneliformis mossae*, revealed nearly 51495 DEGs, of which 11599 were statistically significant (Figure 3b).

**Figure 3.**
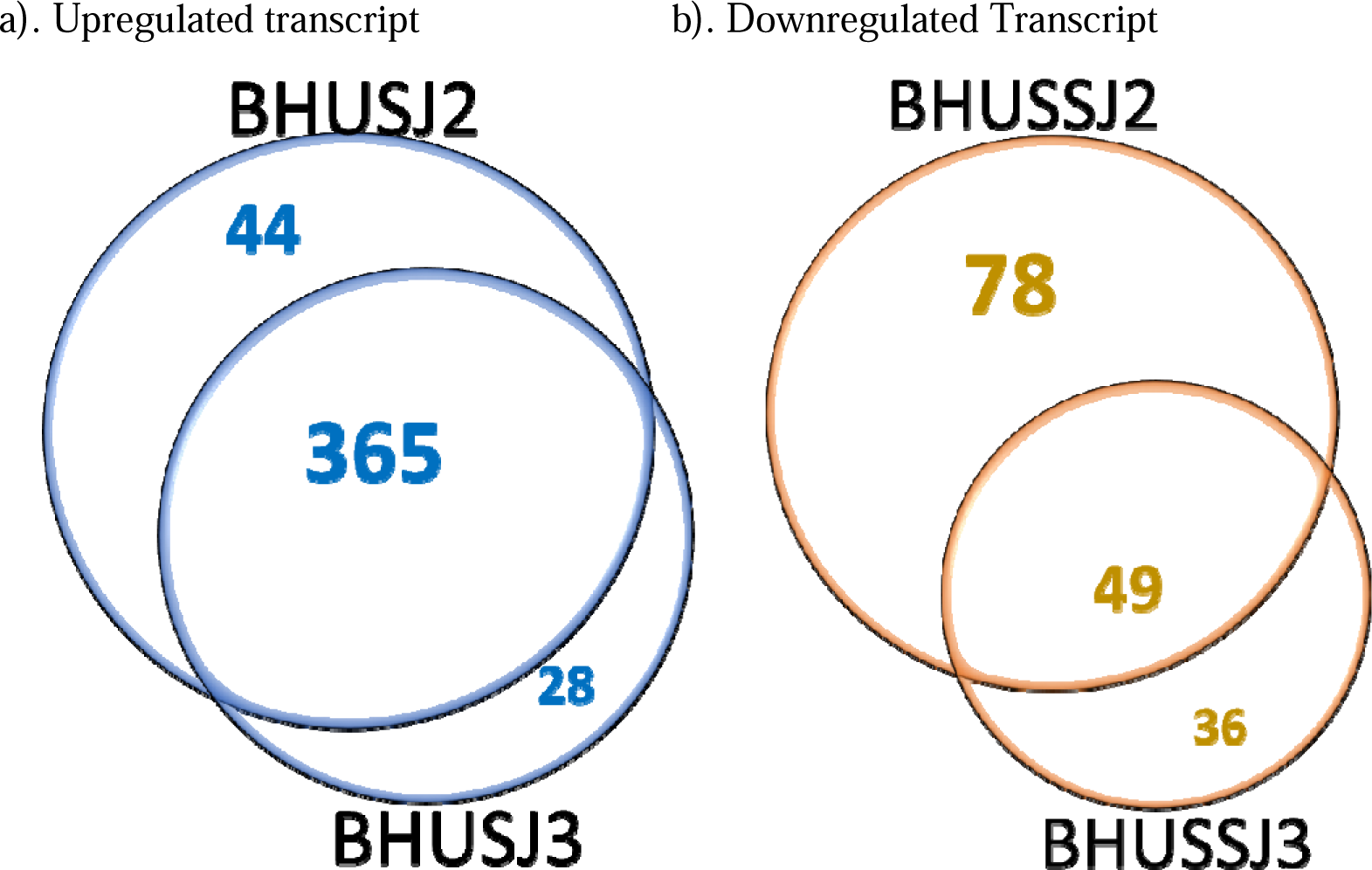
Upregulated and Downregulated transcripts of BHUSSJ2 and BHUSSJ3 in relation to the control samples.

Out of the differentially expressed genes included 4314 upregulated transcripts and 7285 downregulated transcripts. Of the upregulated proteins, 1695 transcripts were upregulated to more than 1.5 folds while among the downregulated transcripts, 575 transcripts were downregulated below −1.5 folds. A total of 365 transcripts were upregulated more than 1.5 folds in *S. melongena* in response to both the fungi, *Glomus macrocarpum* and *Funneliformis mosseae*, while 44 transcripts were upregulated only in response to *G. macrocarpum* and 28 transcripts were upregulated only in response to *F. mosseae* (Figure 4a). Similarly, 49 transcripts were downregulated less than −1.5 folds in response to both the fungi while 78 were downregulated only in response to the *G. macrocarpum* and 36 were downregulated only in response to *F. mosseae* (Figure 4b).

**Figure 4:**
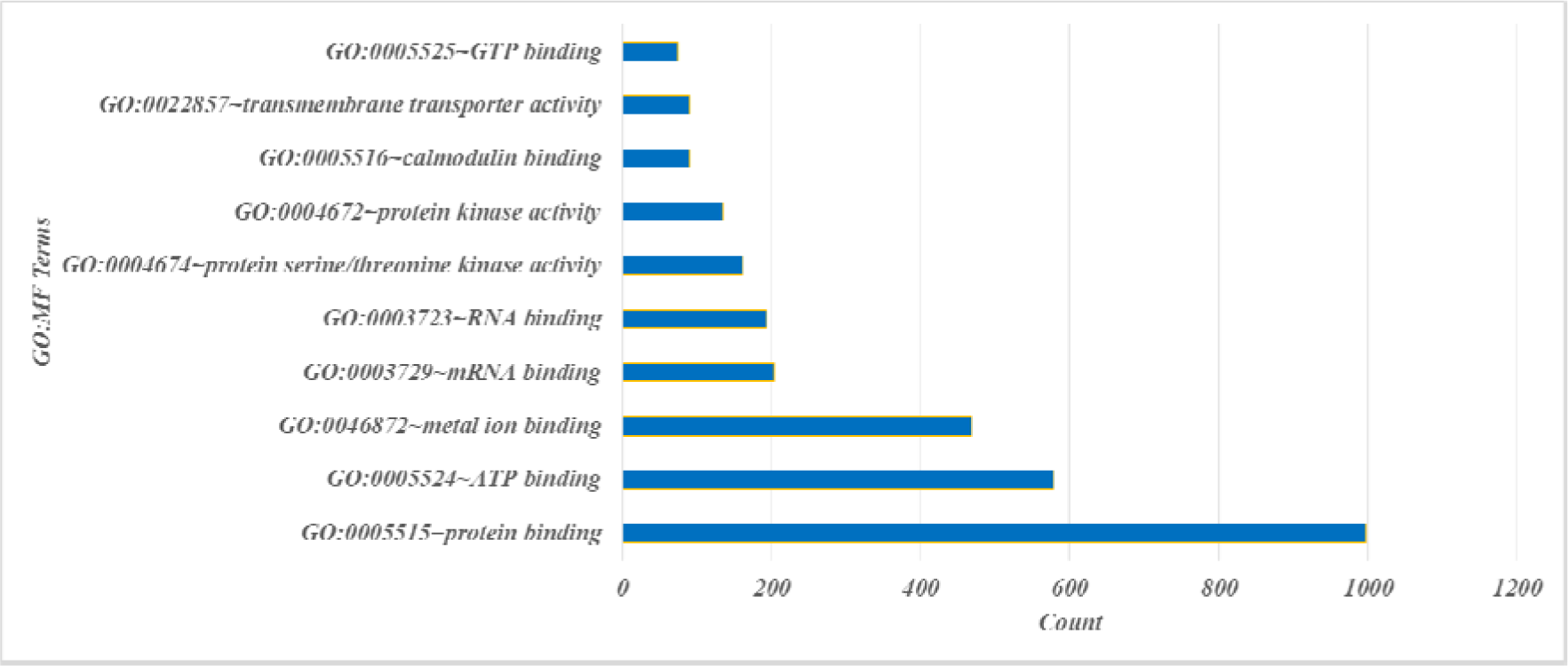
GO annotation representing Top 10 terms in the ‘Molecular Function’ category of Differential expressing transcripts of Control Versus BHUSJ2 Samples

**Figure 5:**
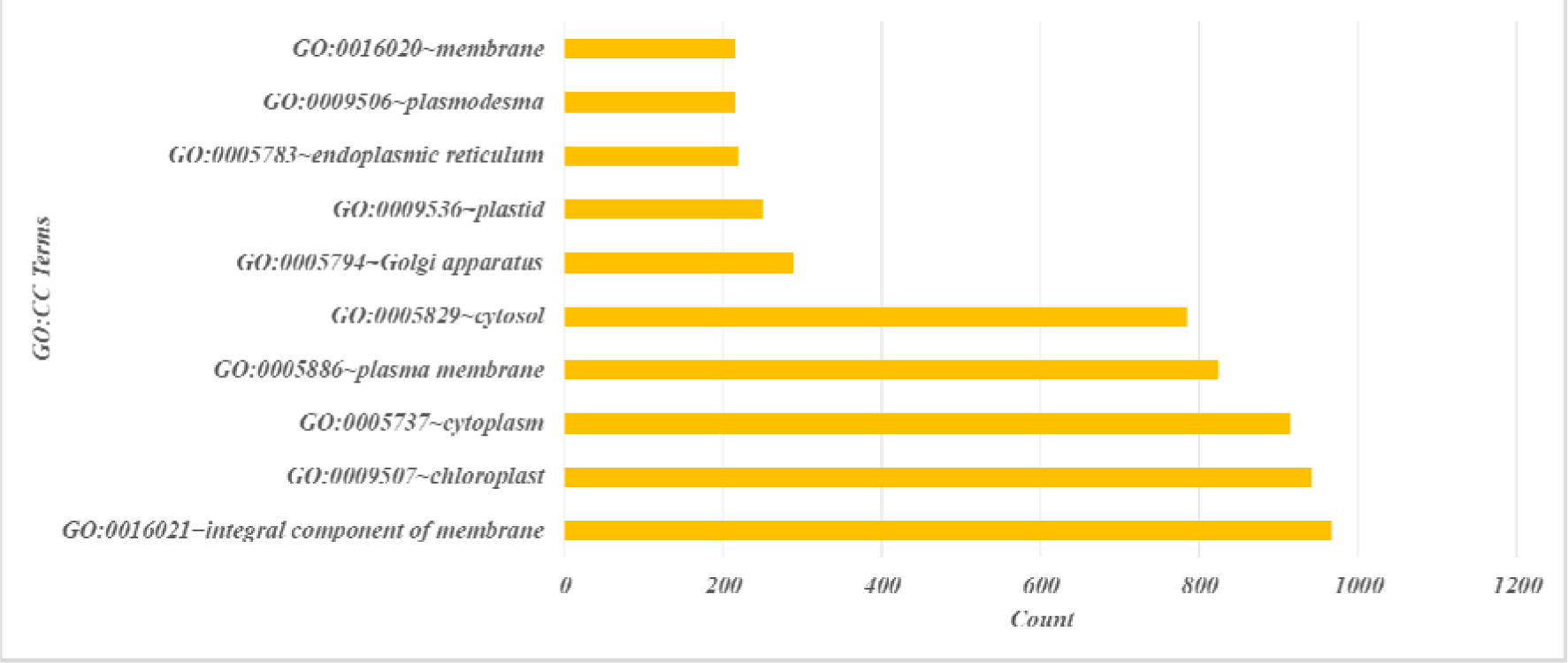
GO annotation representing Top 10 terms in the ‘Cellular Component’ category of Differential expressing transcripts of Control Versus BHUSJ2 Samples

**Figure 6:**
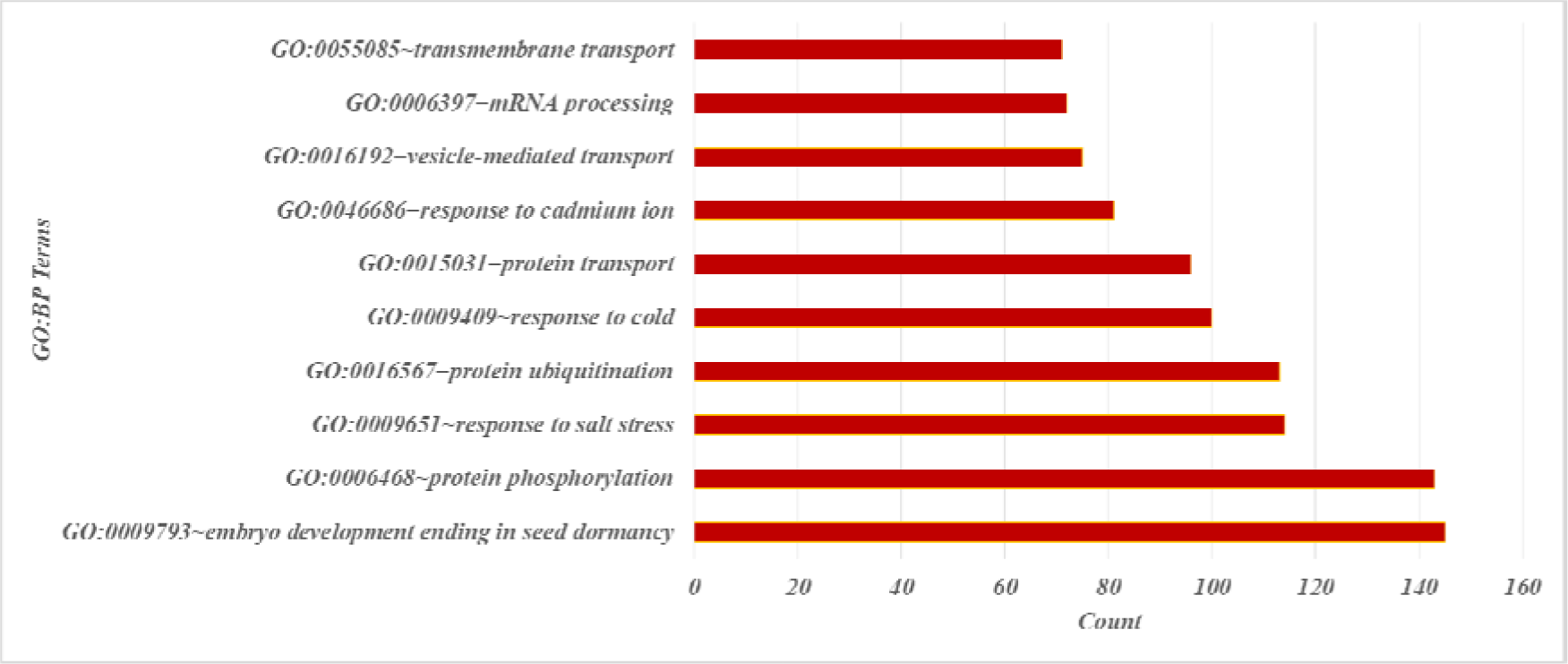
GO annotation representing Top 10 terms in the ‘Biological Process’ category of Differential expressing transcripts of Control Versus BHUSJ2 Samples

### Gene Ontology and KEGG Pathway analysis of the BHUSJ2 and BHUSJ3 samples

The gene ontology analysis by the DAVID software revealed identified 101 different statistically significant “molecular functions” in the differentially expressed genes with the p value less than 0.05. The analysis revealed that the genes belonging to the protein binding dominated the transcript abundance in the BHUSJ2 sample representing the effect of *Glomus macrocarpum* on *S. melongena* with 19% of the genes involved followed by the ATP binding and metal binding activity with the involvement of 11.5 % and 9.3 % of the genes. The analysis of “cellular components” in which the genes were located revealed that the transcript of the membrane, chloroplast and cytoplasm dominated the differentially expressed genes with count 966 (19.2%), 941 (18.67%) and 914 (18.13%) respectively. Similarly, the “biological process” analysis revealed that the differentially expressed transcripts were dominated by the transcript involved in the embryo development leading to breaking the seed dormancy, protein phosphorylation, and salinity responsive transcripts with transcript count 145 (2.93%), 143 (2.89%) and 114 (2.3%) respectively. Pathway KEGG analysis of the DEGs revealed that the top three pathways in which these transcripts were involved included carbon metabolism (76 transcripts - 9.5%), biosynthesis of cofactors (71 transcripts – 8.86%), and Endocytosis (56 transcripts – 7%) (Figure 8)

**Figure 7:**
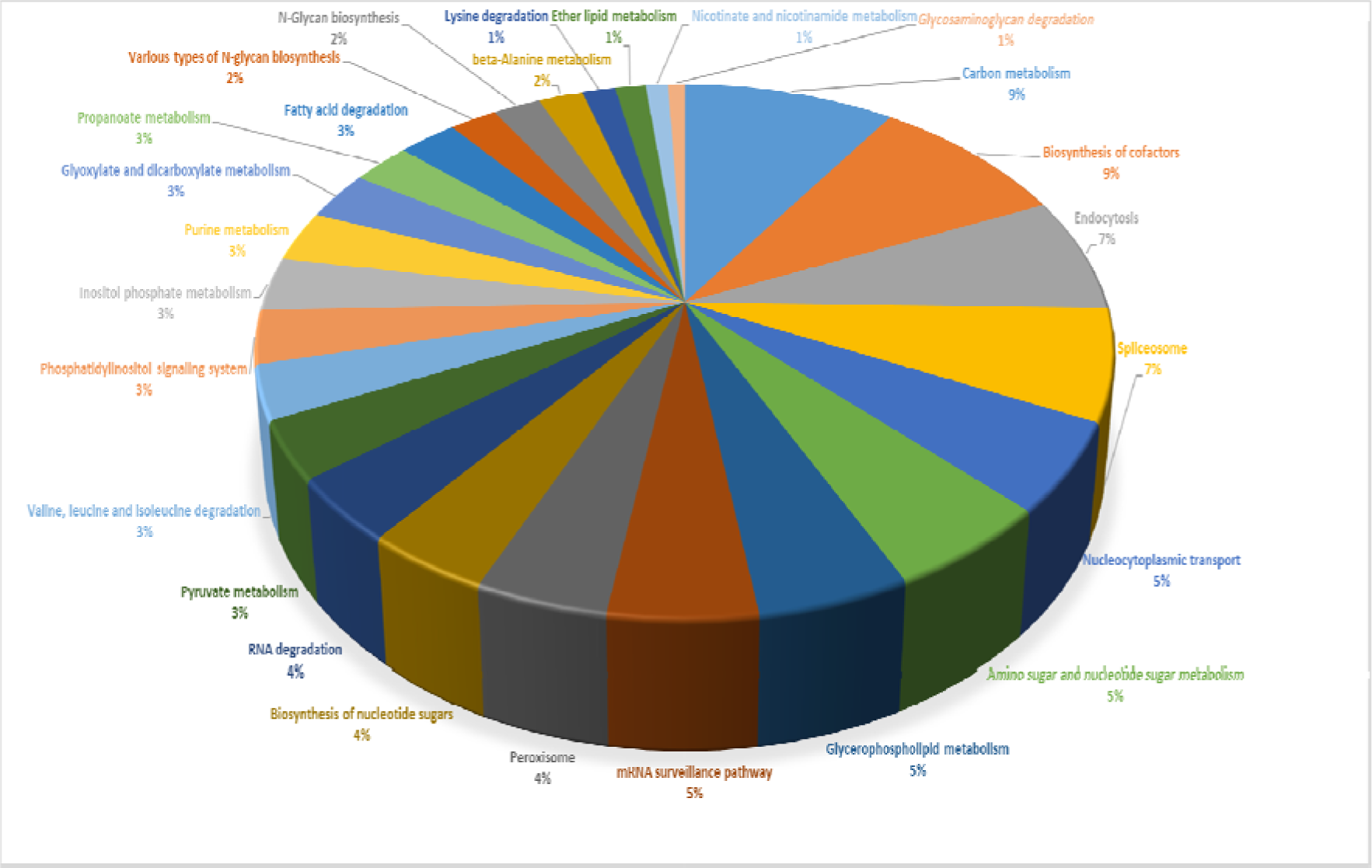
Figure of KEGG analysis representing pathway information related to transcript of Differential expressing transcripts of Control Versus BHUSJ2 Samples

**Figure 8:**
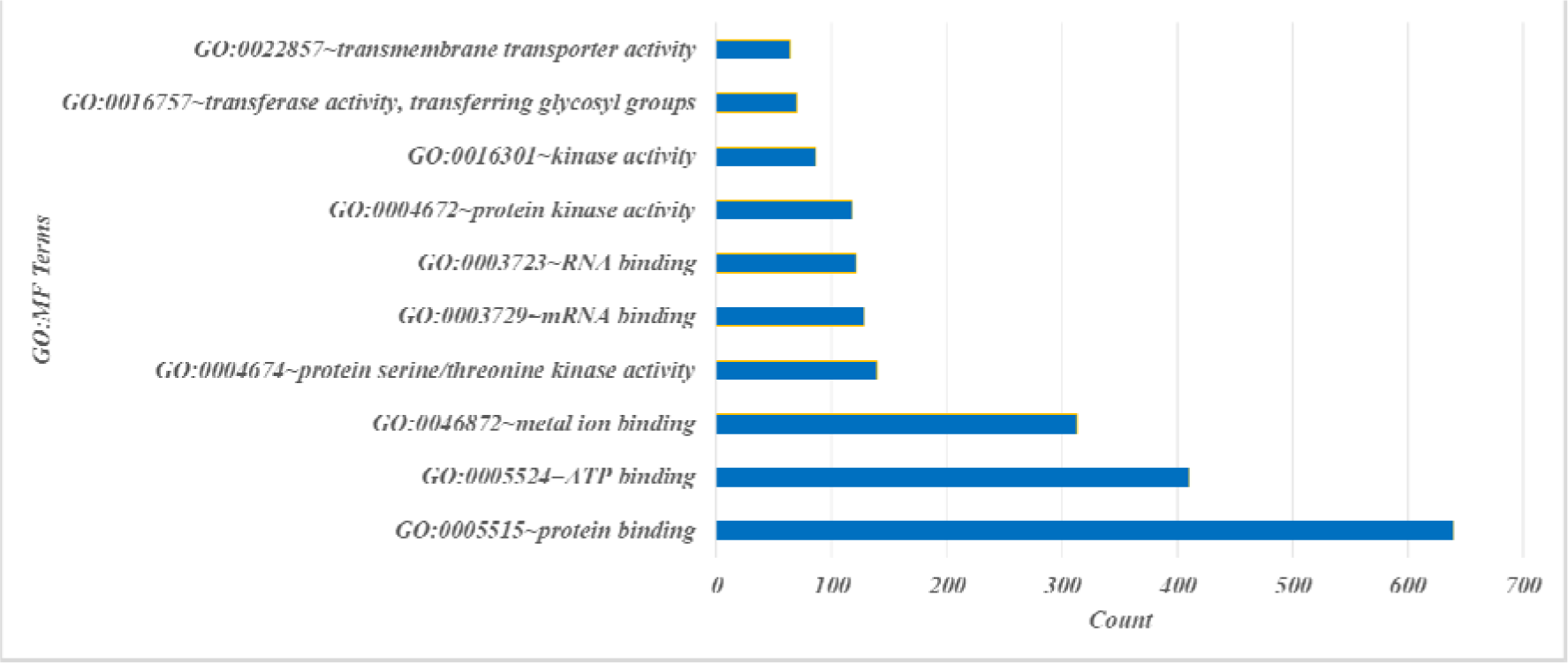
GO annotation representing Top 10 terms in the ‘Molecular Function’ category of Differential expressing transcripts of Control Versus BHUSJ3

**Figure 9:**
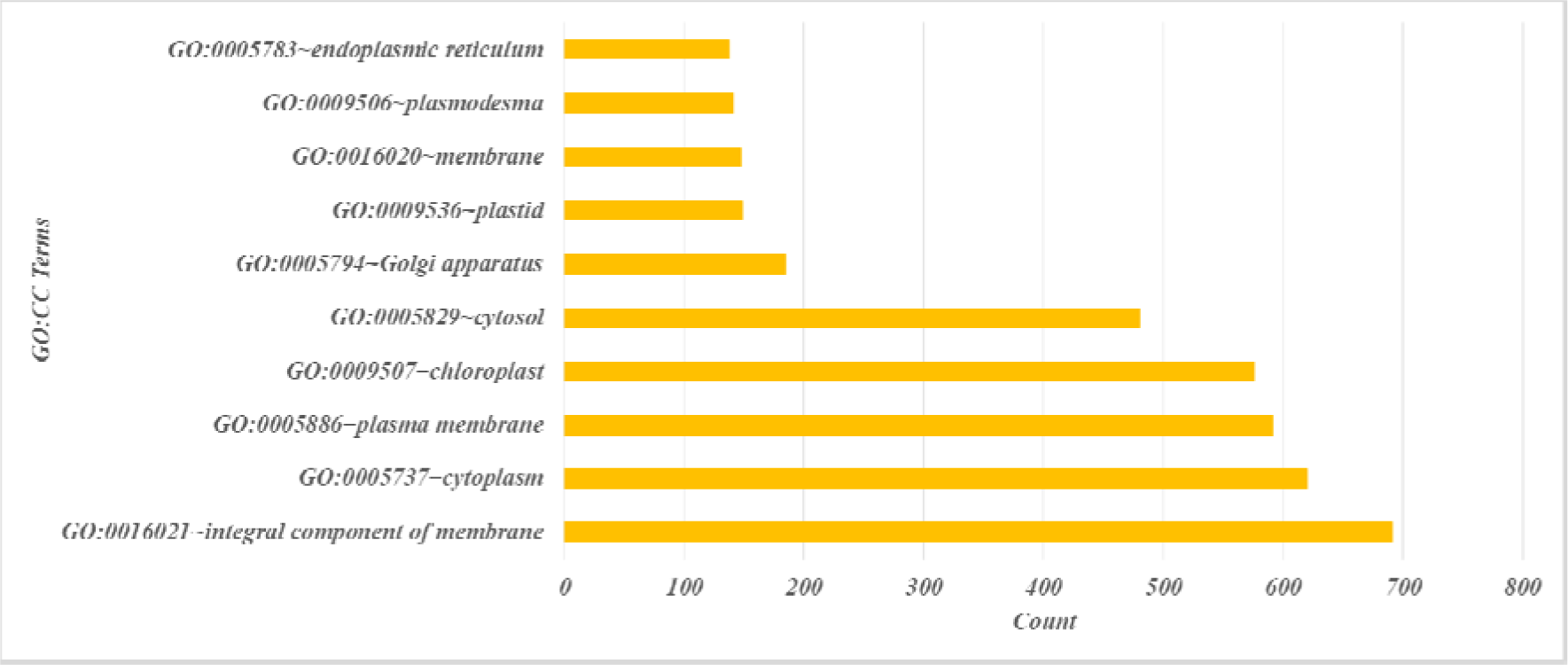
GO annotation representing Top 10 terms in the ‘Cellular Component’ category of Differential expressing transcripts of Control Versus BHUSJ3

**Figure 10:**
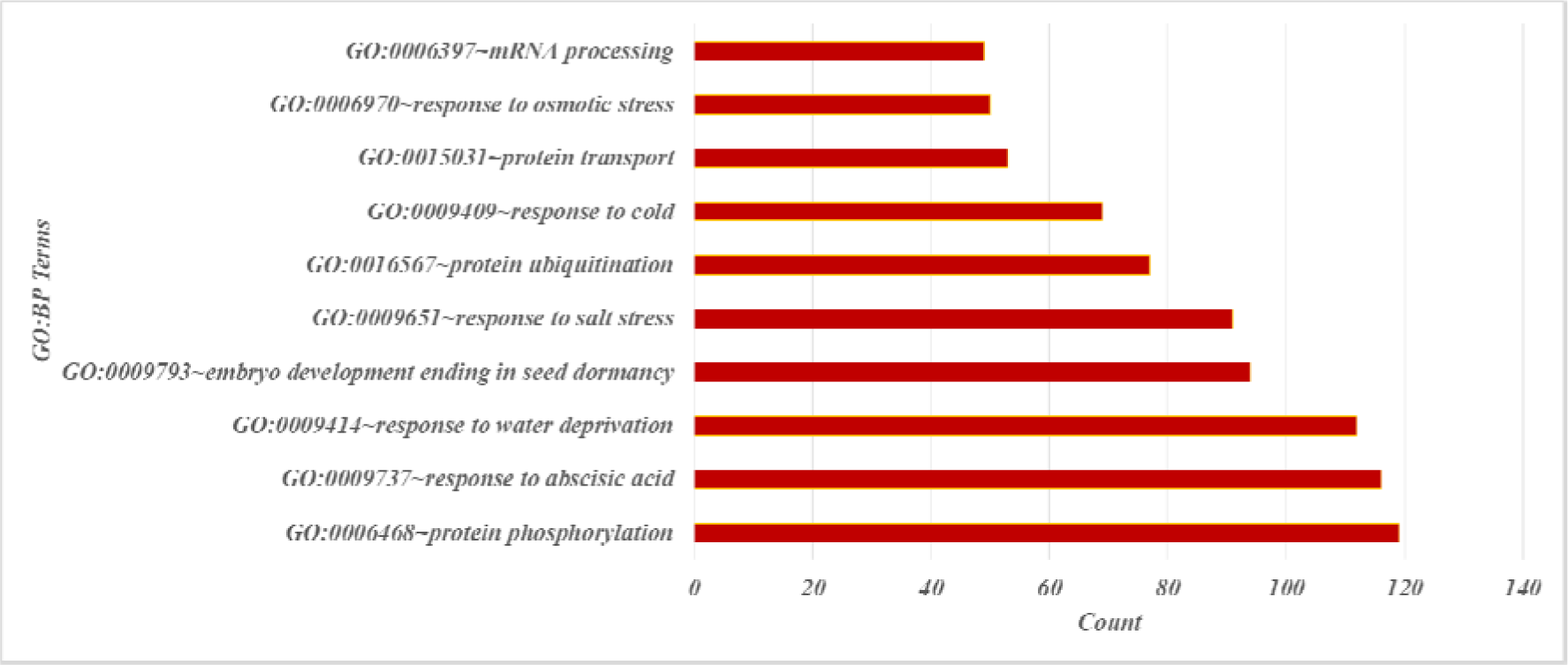
GO annotation representing Top 10 terms in the ‘Biological Process’ category of Differential expressing transcripts of Control Versus BHUSJ3

**Figure 11:**
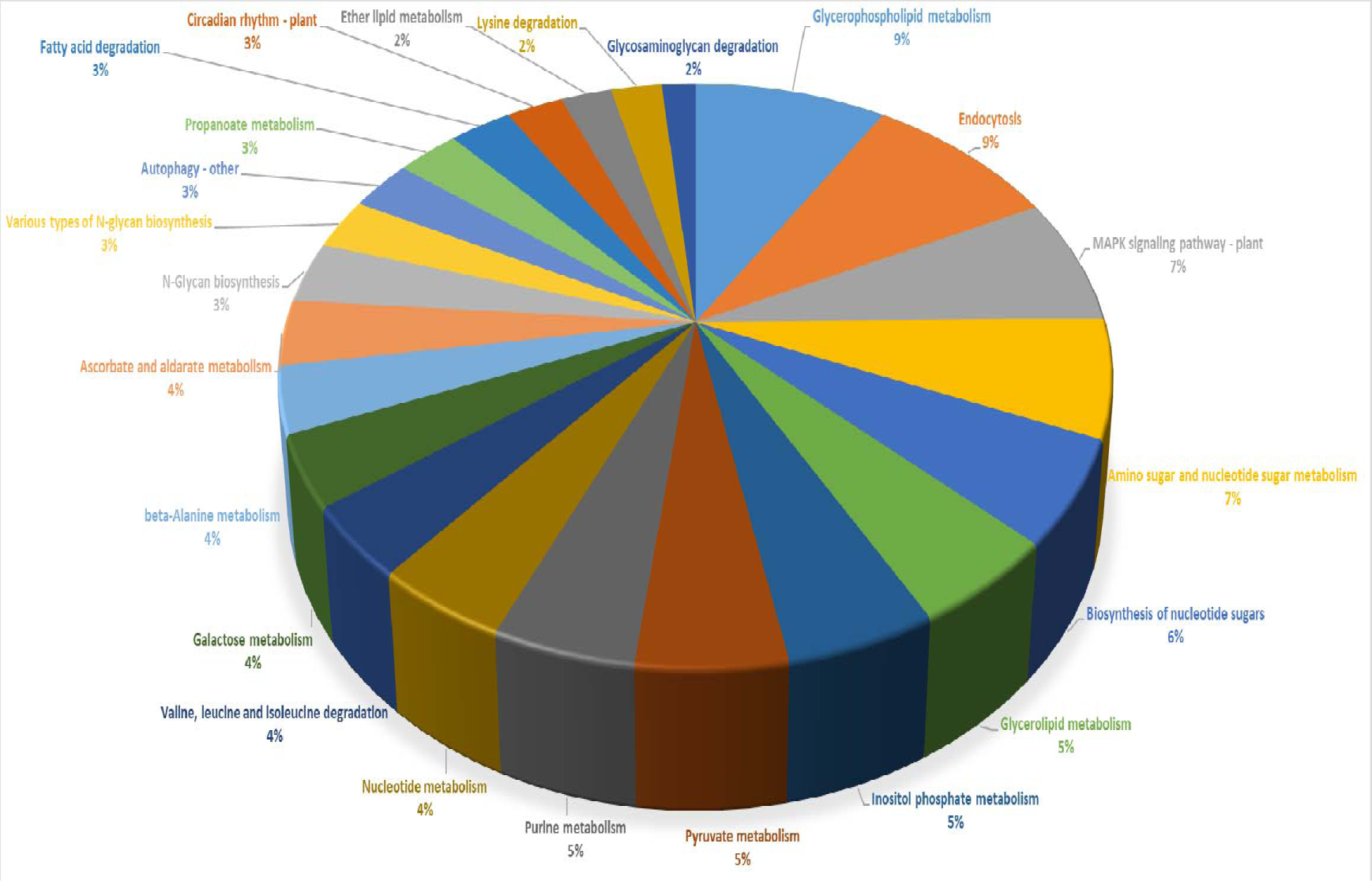
KEGG analysis representing pathway information related to Differential expressing transcripts of Control Versus BHUSJ3.

The gene ontology analysis of BHUSJ3 sample representing the effect of *Funneliformis mosseae* on *S. melongena* revealed that the genes belonging to the function of “protein binding” dominated the DEGs with count = 640 (20.50%) followed by the ATP binding and metal ion binding with count = 410 (13.14%) and 313 (10.02 %) respectively. The analysis of “cellular components” in which the genes were located revealed that the transcript of the membrane, chloroplast and cytoplasm dominated the differentially expressed genes with count 691 (22.14 %), 620 (19.87 %) and 591 (18.94 %) respectively. Similarly, the “biological process” analysis revealed that the differentially expressed transcripts were dominated by the transcript involved in the protein phosphorylation, response to abscisic acid and drought-responsive transcripts with transcript counts 119 (3.81 %), 116 (3.72 %) and 112 (3.6%) respectively. Pathway KEGG analysis of the DEGs revealed that the Glycerophospholipid and Endocytosis metabolism were the dominant metabolic pathway with 34 transcripts (9%) each (Figure 12).

### DEAD-box helicases in the transcriptomes

Among the transcripts involved in the tolerance to salinity stress, DEAD-box helicase motif are known to play significant roles. DEAD-box ATP-dependent RNA helicase 16 (TRINITY_DN6185_c0_g1_i9 and TRINITY_DN2471_c0_g1_i1), helicase 18 (TRINITY_DN15662_c0_g1_i3 and TRINITY_DN20762_c0_g1_i6), and helicase 20 (TRINITY_DN19824_c0_g1_i34) (EC 3.6.4.13) were upregulated to 7.8, 9 and 5.5 folds respectively in *S. melongena* upon association with *G. macrocarpum* and 7.7, 9.7 and 6.9 folds upon association with *F. mosseae*. Another DEAD-box ATP-dependent RNA helicase 18 protein which is also known as, BRI1-KD-interacting protein 115 (BIP115) with Trinity Id TRINITY_DN15662_c0_g1_i5 was upregulated to 2.7x in response to *G. mossae*. Helicases, DEAD-box ATP-dependent RNA helicase 5 (TRINITY_DN22150_c0_g1_i7), helicase 10 (TRINITY_DN26429_c0_g1_i4), helicase 17 (TRINITY_DN8171_c0_g1_i8), helicase 32 (TRINITY_DN48781_c0_g1_i1), helicase 35 (TRINITY_DN1888_c0_g1_i5), helicase 52C (TRINITY_DN1065_c3_g1_i5) and helicase 58, chloroplastic (TRINITY_DN37178_c0_g1_i3) were upregulated to 1.7, 9.9, 2.1, 8.4, 2.7, 3.5, and 8.4 folds respectively in the case of *S. melogena* and *G. macrocarpum* association only. On the other hand, DEAD-box ATP-dependent RNA helicase 22 (TRINITY_DN13923_c0_g1_i1), helicase 36 (TRINITY_DN17609_c4_g1_i5), and helicase 40 (TRINITY_DN14283_c1_g1_i1) were upreguated to 9, 9, and 10.6 folds respectively upon *S. melogena* and *G. macrocarpum* association only. The DEAD-box ATP-dependent RNA helicase 50 (TRINITY_DN37563_c0_g1_i5) was upregulated upon *S. melogena* and *G. macrocarpum* association to 7.9 folds however, another variant of the helicase 50 (TRINITY_DN39418_c0_g1_i3) was decreased to 2 folds in the case of *S. melogena* and *F. mossae* association The modulation of the DEAD box helicases during salinity stress has been correlated with the changes in the superoxide dismutase (Chen, 2016). The levels of superoxide dismutase [Cu-Zn] 2 (EC 1.15.1.1) (TRINITY_DN19272 c1 g1 i4) was upregulated to 9.7x in *S. melongena* in association with *G. macrocarpum* and 7.991x with *F. mosseae*.

**Table 1:**
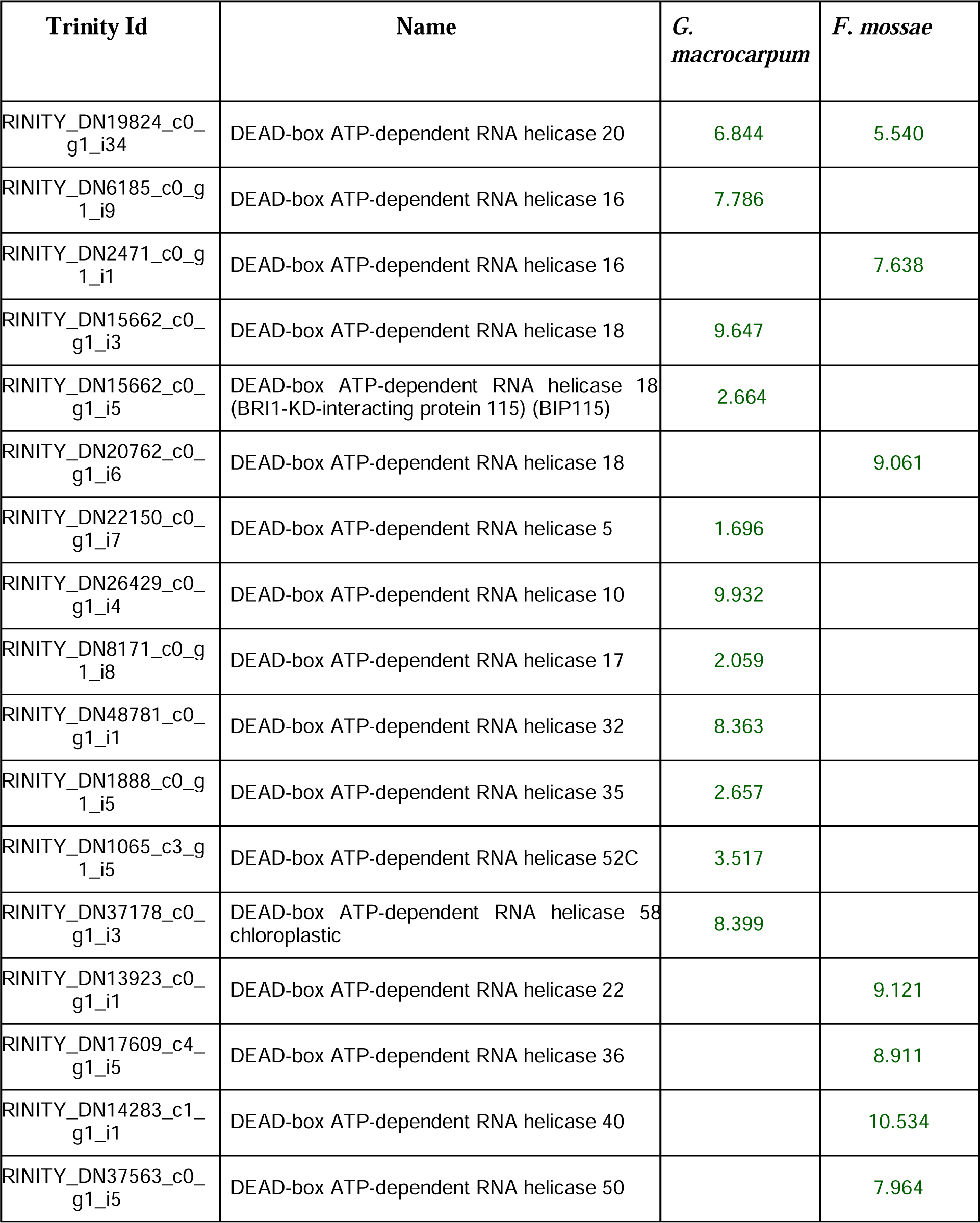
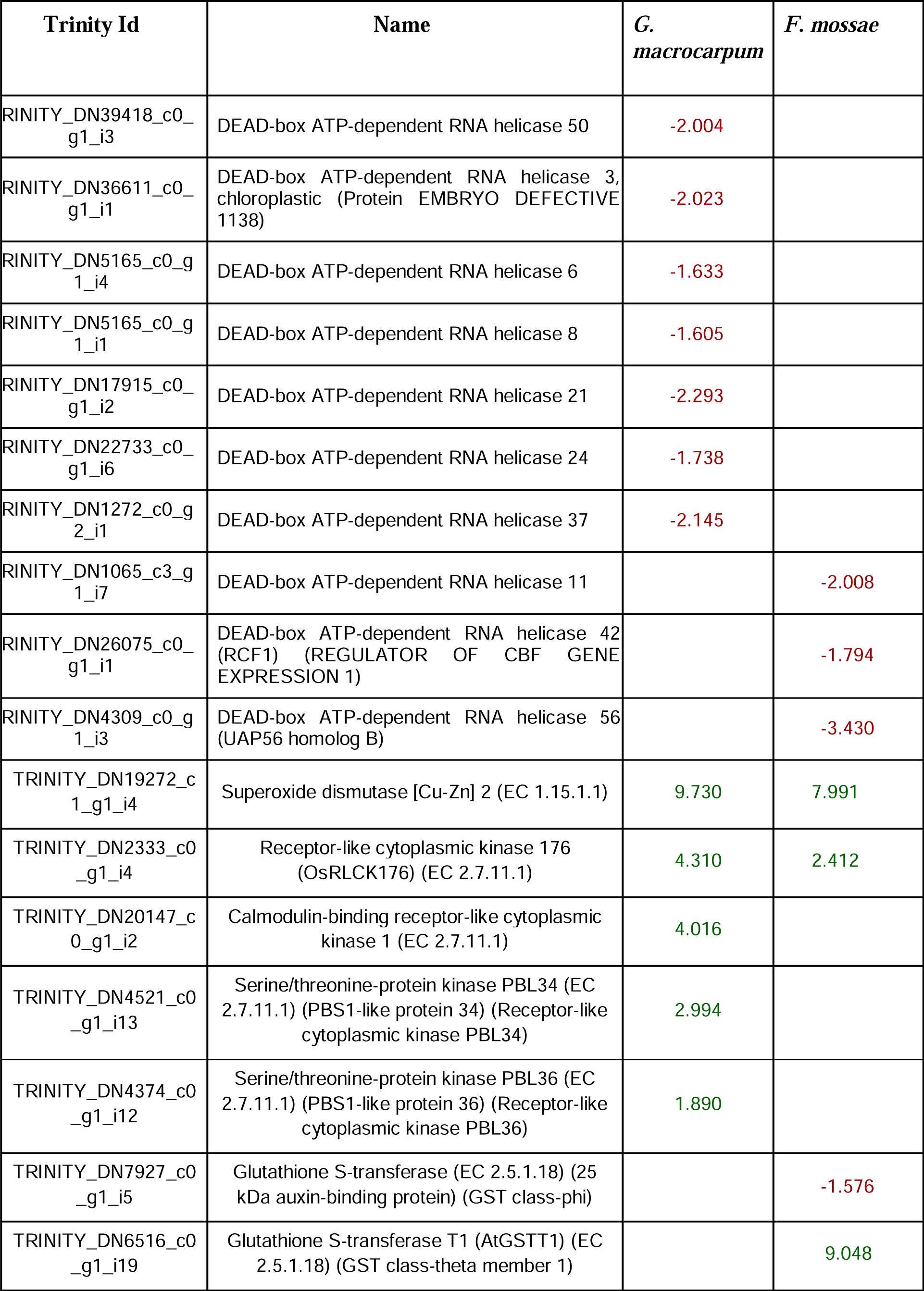
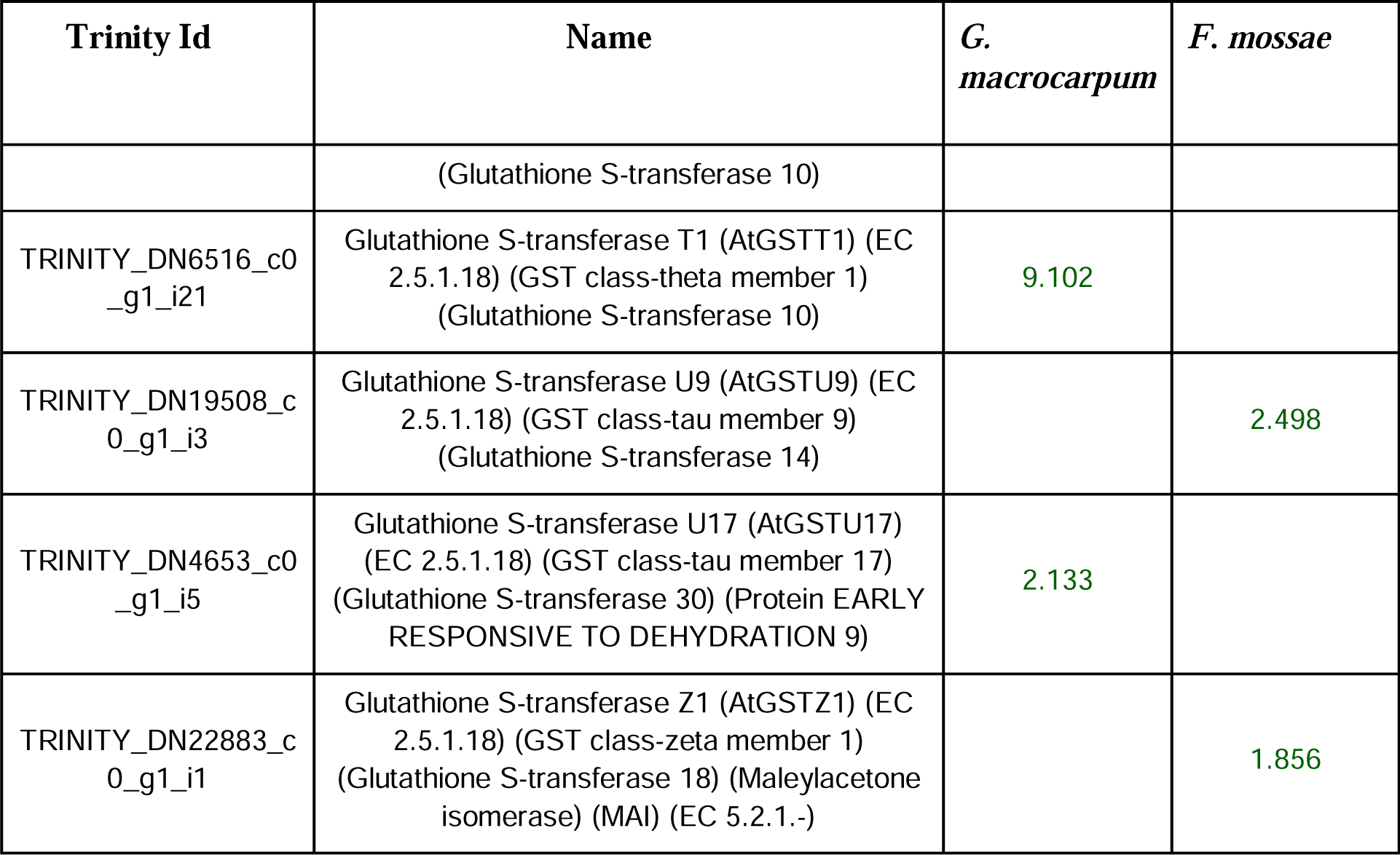
List of significantly modulated DEAD-box ATP-dependent RNA helicases and superoxide dismutase of *Solanum melongena* in association with either *F. mossae* or *G. macrocarpum*.

## Conclusion

Arbuscular mycorrhizal fungi have important role in the agriculture sector. The fungi associate with the roots of plants to form vesicles, arbuscules, and hyphae in roots, and also spores and hyphae in the rhizosphere. The fungi is known to confer a lot of advantage to the host plant in return of shelter including the ability to tolerate stress. The egg plant when treated with two different species of Glomus. viz G. mossae and G. macrocarpum, the two AMF species were found to differentially contribute to the stress tolerance to Solanum melanogena. The findings suggest more studies under salinity stress to gain more insight into the process.

## Acknowledgements

Authors acknowledges UGC for providing the fellowship for this work. We also acknowledge Department of Botany, Banaras Hindu University for providing the necessary lab facilities to carry out these studies.

We do not have any conflict of interest to declare.

## Notes

https://dataview.ncbi.nlm.nih.gov/?archive=bioproject

### Competing Interest Statement

The authors have declared no competing interest.

### Summary of Updates

There are some portions in the material and methods section that i would like to revise.

